# Modeling Transient Brain Coactivity Patterns in Latent Space with FMRI Data

**DOI:** 10.1101/2022.04.28.489899

**Authors:** Kaiming Li, Xiaoping Hu

## Abstract

The brain is a complex dynamic system that constantly evolves. Characterization of the spatiotemporal dynamics of brain activity is fundamental to understanding how brain works. Current studies with functional connectivity and linear models are limited by low temporal resolution and insufficient model capacity. With a generative variational auto encoder (VAE), the present study mapped the high-dimensional transient co-activity patterns (CAPs) of functional magnetic resonance imaging data to a low-dimensional latent representation that followed a multivariate gaussian distribution. We demonstrated with multiple datasets that the VAE model could effectively represent the transient CAPs in the latent space. Transient CAPs from high-intensity and low-intensity values reflected the same functional structure of brain and could be reconstructed from the same distribution in the latent space. With the reconstructed latent time courses, preceding CAPs successful predicted the following transient CAP with a long short-term memory recurrent neural network. Our methods provide a new avenue to characterize the brain’s transient co-activity maps and model the complex dynamics between them in a framewise manner.

## INTRODUCTION

The brain is a complex dynamic system with multiple networks interacting with each other in a dynamic and coordinated manner (Shine et al 2019, Spadone et al 2015). Characterization of the complex spatiotemporal dynamics is fundamental to advancing our knowledge of how the brain works. With resting state functional magnetic resonance imaging (rs-fMRI), recent studies have begun to investigate temporal patterns by time-varying functional connectivity (FC) analysis (Calhoun et al 2014, Chang & Glover 2010, Hutchison et al 2013, Sakoglu et al 2010). However, many prior efforts used a sliding window strategy that calculated a consecutive series of FC matrices, each of which was defined over a short period of a fMRI scan, e.g., 40 seconds (Allen et al 2014, Aruljothi et al 2020, Vanni et al 2017, Zalesky & Breakspear 2015). While this type of analysis may capture some low-frequency dynamics, its sensitivity in rapid dynamic change detection is limited. Furthermore, FC and other methods previously reported (Beckmann et al 2005, Taghia et al 2018, Vidaurre et al 2017) are essentially linear, which may be insufficient in modeling the brain’s complex spatiotemporal dynamics.

There have been multiple efforts on characterization of fMRI brain activity in a framewise manner. Liu and Duyn (Liu & Duyn 2013) examined the co-activation patterns of brain at different time frames of the fMRI scan. They found that high-intensity brain voxels at a given time frame resembled brain networks, or part of them, obtained from the FC and independent component analysis, providing important insights on how brain networks evolve and interact with each other over time. Karahanoğlu et al. decomposed BOLDs for the activity-inducing driver using rsfMRI data (Karahanoglu et al 2013). Based on the resultant activity-inducing signals, not only the external stimuli or spontaneous activity of brain regions were captured, the non-stationary relationships between different networks were also revealed. While these framewise methods enable the examination of spatial patterns at the highest possible temporal resolution for fMRI data, the temporal information between frames is not explicitly modeled, and the temporal features can only be examined in an ad-hoc manner.

The past decade has witnessed remarkable development of deep learning techniques in computer vision and their applications in brain imaging studies (Goodfellow et al 2016). For spatial information representation, variational autoencoder (VAE) (Kingma & Welling 2013) can build complex generative models and map complex spatial patterns to a latent space following a multivariate gaussian distribution. The introduction of stochasticity makes the model robust to small changes in the latent variables. VAE has been used to explain and predict fMRI responses to natural videos (Han et al 2019), reconstruct faces from fMRI patterns (VanRullen & Reddy 2019) and find connectivity states with rs-fMRI (Qiang et al 2020, Zhao et al 2019). For temporal modeling, a variety of recurrent neural networks (RNN), including gated recurrent unit (Cho et al 2014) and long short-term memory (LSTM) RNN (Hochreiter & Schmidhuber 1997), have been successfully developed for dynamic analysis beyond linearity(Guclu & van Gerven 2017) and even for predicting future activity based on prior data (Sobczak et al 2021). RNN was also able to accurately identify individuals based on short clips of fMRI data, as it captured the complex spatiotemporal features from rs-fMRI data instead of merely spatial patterns in the connectome (Chen & Hu 2018, Wang et al 2019). These neural networks provide us powerful tools to represent the spatial patterns and model them temporally.

Inspired by the concept of transient co-activation maps (Liu & Duyn 2013), the present study mapped the high-dimensional transient co-activity maps of fMRI data in a low-dimensional latent space with a generative variational auto encoder (Kingma & Welling 2013). We evaluated the performance of VAE on multiple datasets at different resolutions and with both high-intensity (or positive-value in a normalized time course) and low-intensity (negative-value) of fMRI frames. With the reconstructed latent time courses, we examined whether prior CAPs could predict the following transient CAP using a LSTM recurrent neural network. Our study provides a new way to characterize brain’s transient co-activity maps and model the complex dynamics between them in a framewise manner.

## MATERIALS AND METHODS

### Datasets and preprocessing

Three public brain imaging datasets (ABIDE-1 (Di Martino et al 2014), HCP (Van Essen et al 2013) and B-SNIP (Tamminga et al 2014)) were used in the present study. Detailed description of the study design, ethical approval and subject recruitments has been provided by each project (Di Martino et al 2014, Tamminga et al 2014, Van Essen et al 2013). For the ABIDE-1 and B-SNIP datasets, resting-state fMRI data were preprocessed with a customized script primarily based on AFNI (ver. 16.2.07) (Li et al 2020). Specific preprocessing steps included despiking, slice time correction, motion correction, rigid registration onto the high-resolution T1 image, and smoothing with an FWHM of 4mm and nuisance regression (regressors: 3 translations, 3 rotations, 1^st^ order derivatives of the 6 registration parameters and 3 principal components of CSF signals). The residual signals were further cleaned using ANATICOR(Jo et al 2010), which modeled and removed local transient hardware artifacts. Finally, the processed BOLD images were affinely registered onto the 3 mm isotropic MNI152 standard space using the T1 image as an intermediate target. For the HCP dataset, we used the preprocessed dataset directly.

### Transient co-activity maps

After the above preprocessing, the time course of each gray matter voxel was z-scored and thresholded to obtain the *activated* (positive values in a normalized time course) and *deactivated* (negative values in a normalized time course) time frames in the recorded fMRI session. Two thresholds, i.e., the 85^th^ (Liu & Duyn 2013), and the 15^th^ percentiles of the examined time course, were applied respectively, with the 85^th^ percentile for the activated frames (zeroing any frames below it) and the 15^th^ for the deactivated time frames (zeroing any frames above it). Each thresholding operation was performed temporally based on the brain activity of the same brain region, providing normalization for regions across the brain.

Spatially, at each time frame, each thresholding resulted in a transient CAP, either co-activated or co-deactivated. The resultant transient CAPs were then spatially downsampled to 1 × 1 × 1 cm^3^ for subsequent analysis. Note that the negative values resulted from the 15^th^ percentile thresholding were flipped for visualization and comparison purposes.

### VAE model

The VAE model, as shown in Figure 1A, contains an encoder and a decoder. The encoder maps an overt high-dimensional transient CAP image *x* in the image space to a lowdimensional variable *z* in the latent space. Considering the high individual variability of subjects in brain anatomy, the encoder represents a statistical distribution of individual transient CAP *x: q*_ø_(*z/x*). Given a variable *z* in the latent space, the VAE model will generate a sample *x* with the decoder, which is a statistical model *p*_ø_(*x/z*). Note that both *x* and *z* are multivariate vectors.

**Figure 1.**
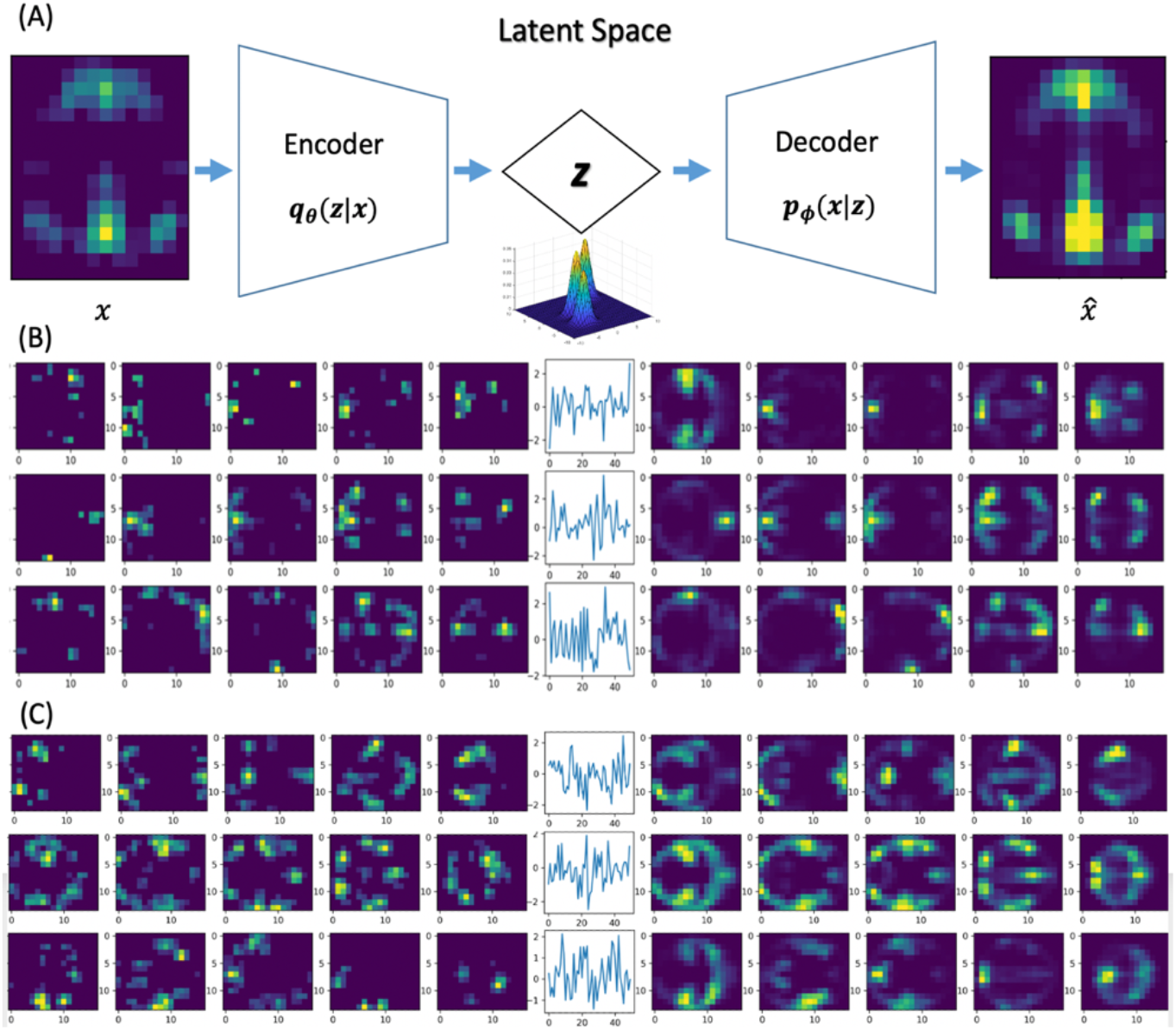
(A) a scheme of the VAE model. (B) raw transient CAPs, latent variables, and reconstructed CAPs using the ABIDE-1 dataset. Each time series describes a latent variable z (N=50). The five slices on its left show a raw transient CAP or *x* while the five slices on its right depict a reconstructed sample 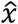. For each slice, top is left hemisphere, and bottom is right hemisphere. The tested images are from the ABIDE-1 dataset. (C). Reconstruction of B-SNIP CAPs with the model trained on ABIDE-1.

The variable z follows a multivariate gaussian distribution. It can be parametrized by the distribution mean μ and variance σ or log *σ* via a random variable *ε: z* = *μ* + *σ*⨀*ε*, where *ε* follows the normal distribution *N*(0,1).

Specifically, after reshaping the 3D CAP into 1D and cropping the non-brain background, each transient CAP image *x* was represented as a vector of 3332 image features. A single hidden layer, with 2222 intermediate neurons as suggested earlier (Heaton 2008), was used to map the input features to the hidden space. Then, two separate layers were used to train the mean μ and the log variance log *σ* of the distribution. The dimension of *z* was set to 30 for low-resolution datasets (ABIDE-1 and B-SNIP) and to 50 for high-resolution datasets (HCP) or for comparison across datasets.

Training of the VAE model was performed on the FastAI platform (Howard & Gugger 2020). The loss function was defined as the combination of a reconstruction loss, which was the binary cross entropy between the raw and the reconstructed images, and a regularization loss, which was the Kullback-Leibler divergence of latent distributions. The weight parameters were initialized using He’s method (He et al 2015), and optimized using the Adam algorithm (Kingma & Ba 2014). To prevent overfitting, dropout layers were added after the rectified linear unit activation layers of VAE. For each dataset, the subjects were randomly split into a training dataset and a validation dataset with a sample size ratio of 7:3. To reduce noise, volumes with fewer than 30 remaining voxels after thresholding were dropped.

### Visualization of the latent dimensions

After training, we generated a sample image for a unit latent vector along each latent dimension using the resultant decoder and visualized them in the image space. To illustrate the relationship amongst these latent unit vectors, a hierarchical clustering algorithm was performed to aggregate them based on the similarity of their reconstructed sample images. Euclidean distance was used as the similarity metric. Due to the limit of page layout, we only showed the result of 30 latent dimensions obtained on the ABIDE-1 dataset.

### Comparison of models trained with positive and negative values

The images thresholded by the 85^th^ and the 15^th^ percentiles at a given time frame correspond to the activated (positive-value) and deactivated (negative-value) brain regions at that time, respectively. To examine whether the positive and negative values carry similar information regarding the brain’s functional organization, two VAE models were trained using the positive and negative values from the same dataset, respectively. The trained models were compared on their latent structures and their ability to reconstruct transient CAP_s_.

Specifically, two rs-fMRI sessions, i.e., rfMRI_REST1_LR or R1LR and rfMRI_REST2_LR or R2LR(Van Essen et al 2013), were used in this analysis. The positive and negative values from R1LR were used to train the two VAE models and those from R2LR were used as testing. The parameters of both models were plotted as matrices for qualitative comparison. The derived latent variables were similarly visualized in the image space as above. The resultant images from the two models were matched so that the mutual information between each pair was maximized. The reconstruction losses of both models in the separate dataset R2LR were statistically compared using the paired student t-test.

### Prediction of transient CAPs with reconstructed latent time courses

To examine whether the mapping of image features to the latent space were effective and whether they captured the underlying brain dynamics, we applied an LSTM recurrent neural network on the reconstructed latent time courses of the R1LR dataset to predict the subsequent transient CAP with preceding ones.

To do so, the transient CAPs were first mapped to the latent variables with the trained VAE encoders. The latent variables were concatenated over frames and formed latent time courses. The resultant time courses, with the same number of frames as raw fMRI signals, were then broken down into 40-frame segments. These segments were then fed into LSTM to predict the next latent variable that corresponded to the 41 ^st^ time frame.

The LSTM model we used was based on AWD-LSTM (Merity et al 2017), where dropout, activation regularization and temporal activation regularization are combined to reduce overfitting in the training process. We also tested the performance of stacked and bidirectional AWD-LSTM for the prediction.

## RESULTS

### Transient CAPs mapping and reconstruction

Figure 1A demonstrates the scheme of our VAE model. The left side shows a transient CAP *x* where the default mode network was activated. With the encoder, *x* was mapped to a latent variable *z* that followed a multivariate Gaussian distribution in the latent space. With the decoder, *z* was successfully reconstructed in the image space as a transient CAP 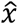. Qualitatively, the reconstructed CAP 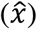 highly resembled the raw transient CAP (*x*).

Panel 1B illustrated three random cases of mapping and reconstruction using the VAE model trained on ABIDE-1 dataset. Each row depicts one case. From the left to the right are the original transient CAP *x*, the encoded latent variable *z* and the reconstructed transient CAP 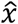, respectively. As seen, the VAE model not only captured main bodies of the original CAPs and but also denoised them.

In particular, with the VAE model trained on the ABIDE-1 dataset, we were able to reconstruct the transient CAPs in the B-SNIP dataset (Figure 1C), suggesting that this model is robust across datasets with similar image resolutions.

### Representation of latent dimensions in image space

We generated images for the unit latent vectors along each dimension using the trained decoder and visualized them in Figure 2. Figure 2A plots the corresponding dendrogram of the hierarchical clustering. Each number on the y-axis corresponds to the image representation of a latent dimension. There are 30 latent dimensions shown in total. The resultant images (Figure 2B) were arranged in the same order as in Figure 2A.

**Figure 2.**
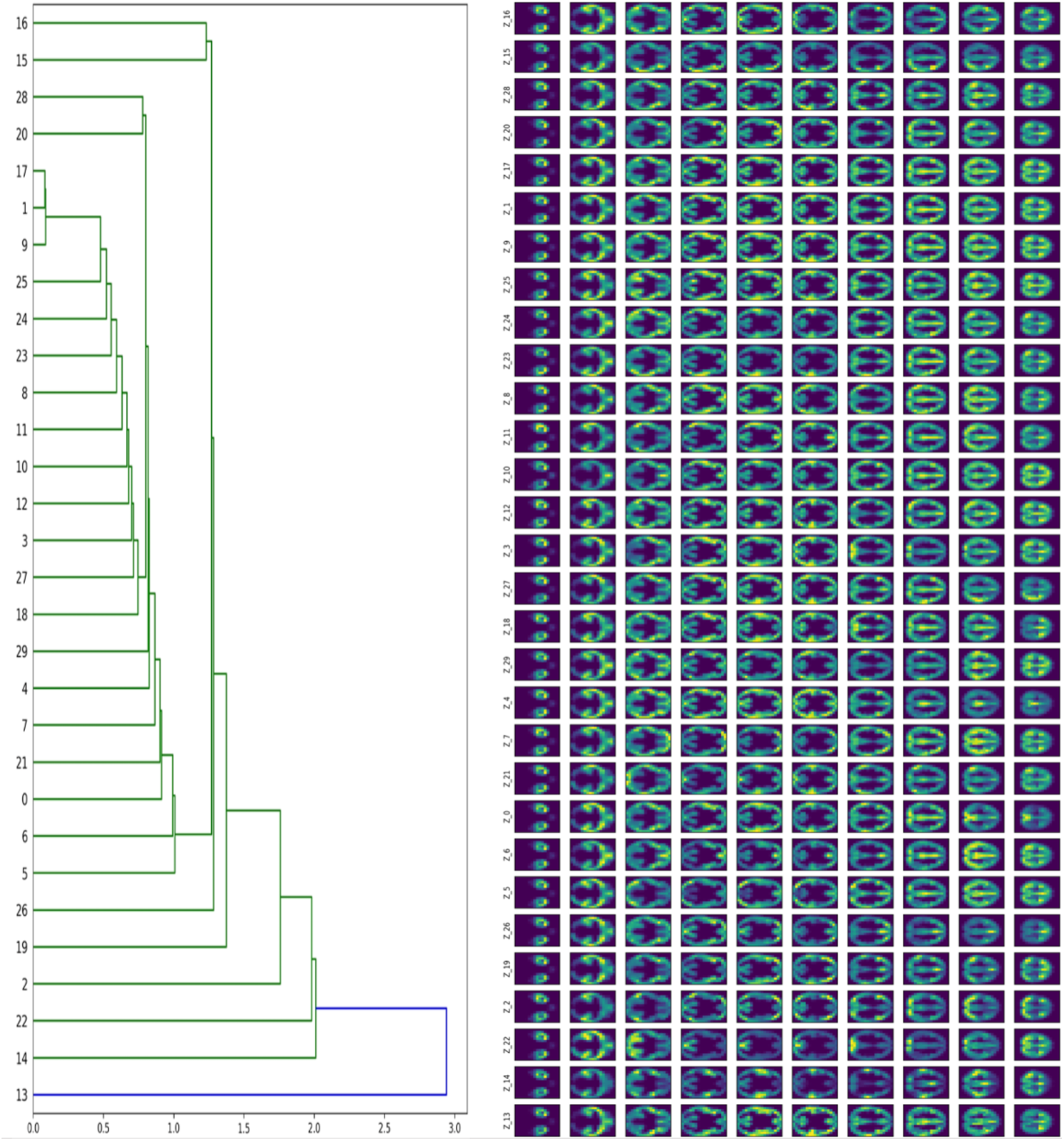
Left: Hierarchical clustering of the representations of latent dimensions (*N*=30) in image space. Right: the corresponding image representations of latent dimensions. The model was trained on ABIDE-1.

As seen from the images in Figure 2B, the gradually changing patterns resemble the resting state networks obtained from the traditional FC analysis and independent component analysis.

Although there are overlaps in the gray matter areas, especially for the neighboring images, there are significant differences in the prominent activated regions between them.

### Models trained with low-/high-resolution datasets

We examined the VAE models trained with low-resolution datasets (ABIDE-1 and B-SNIP, i.e., ABIDE+BS) and high-resolution ones (the four HCP rs-fMRI sessions, i.e., REST1_LR, REST1_RL, REST2_LR, and REST2_RL). The results are summarized in Figure 3. Since there may be no correspondences in the inner layers of the VAE models, we only visualized and compared the first layer (FC1, the upper panel in Figure 3) and the last layer (FC4, the lower panel in Figure 3) of the derived VAE models. Essentially, FC1 maps the image features into the hidden layer in the encoder, while FC4 maps the hidden features to the image features in the decoder.

**Figure 3.**
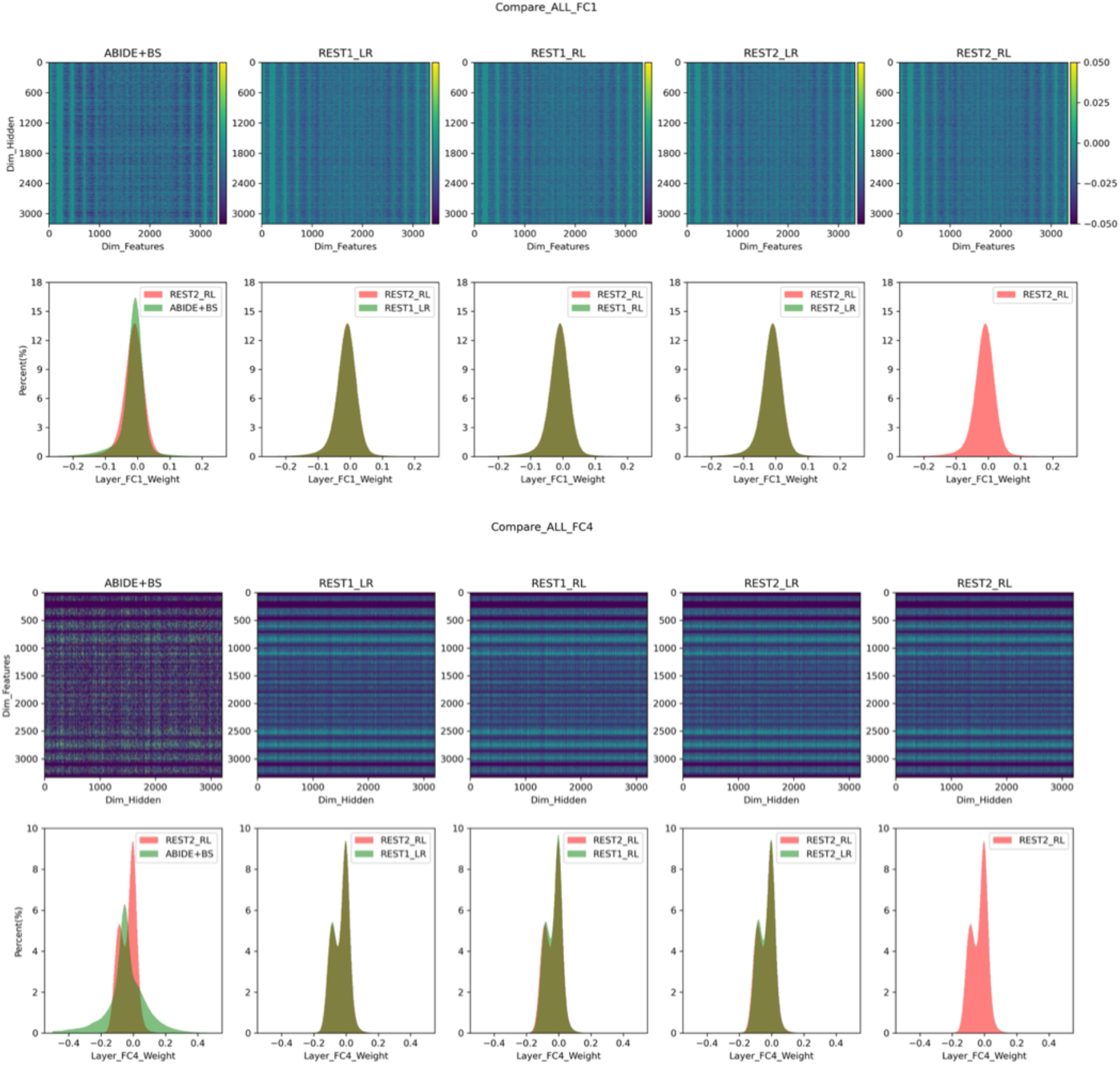
Comparison of VAE models trained on different datasets. ABIDE+BS is the low-resolution ABIDE-1 and B-SNIP datasets, while REST1-LR, REST1-RL, REST2-LR, and REST2-RL are high-resolution HCP rs-fMRI datasets. The first layer FC1 and the last layer FC4 were depicted. Each showed the weight matrices (upper row) and the weight distributions (lower row). Note that all models were initialized with the model trained on REST2_RL.

For the four HCP rs-fMRI sessions, the trained models were very similar for both FC1 and FC4, as seen in both the weight matrices (upper row) and the weight histogram (lower row). The model from the low-resolution datasets was generally like those from the HCP sessions but had lost details as revealed by the sparser weight matrices, particularly in the last layer FC4. We further examined the differences in FC4 between low- and high-resolution models and found that they mainly resided in the surroundings of brain, the temporal lobe, and regions close to the sinus (Figure 4). These regions usually suffer signal loss and have image distortion in fMRI acquisition.

**Figure 4.**
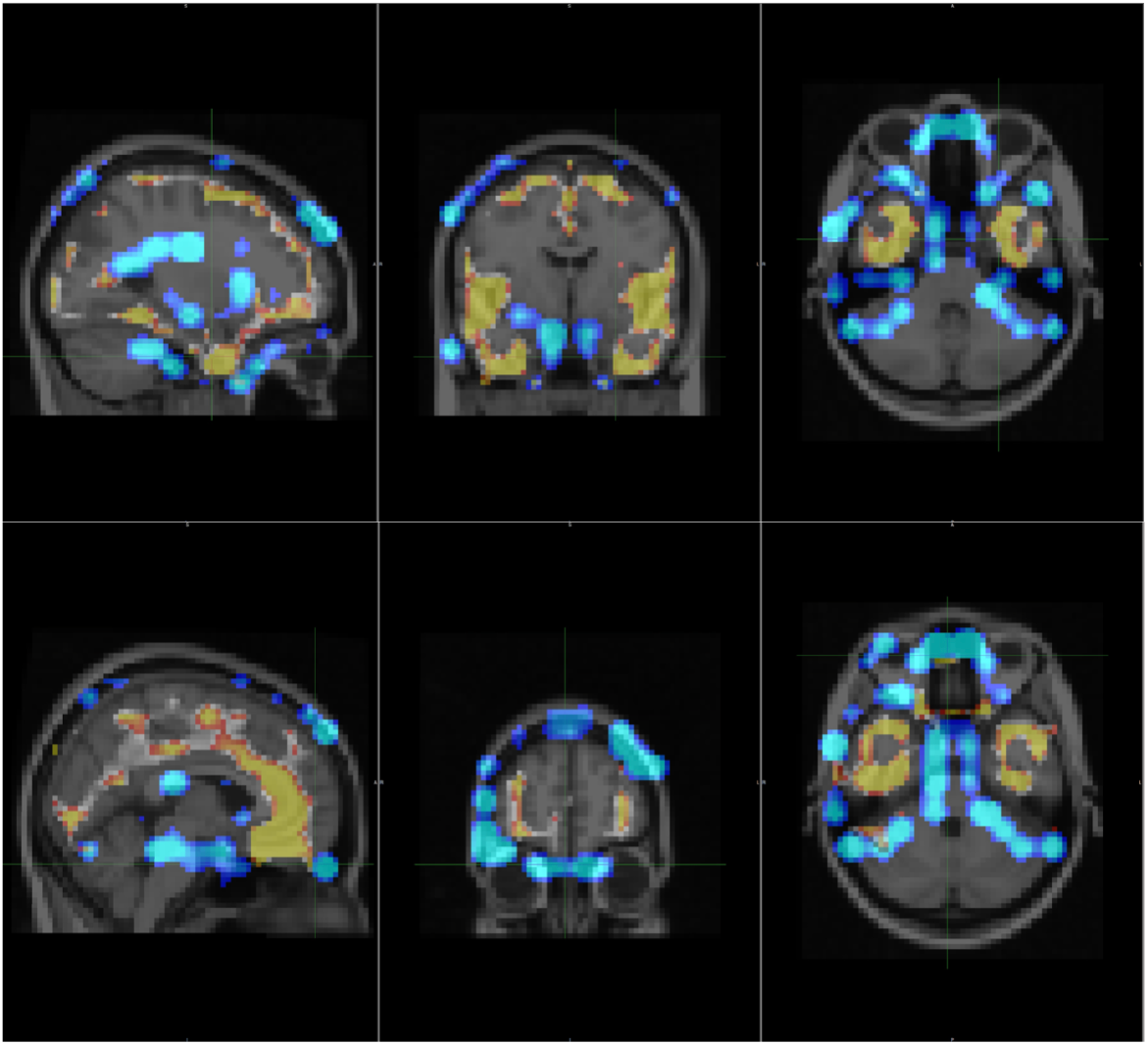
The differences in layer FC4 between low- and high-resolution models. Warm color regions affect the low-resolution model, while cold color regions affect the high-resolution model.

### Models trained with positive values and negative values

The model differences in reconstruction losses are depicted in the panels B and C of Figure 5, respectively. For the positive-value (PosVal) dataset of R2LR, the PosVal model had a superior performance compared to the negative-value (NegVal) model, and vice versa. However, the performance gains, defined as the extra loss reduction acquired by running a model on its corresponding native dataset, were 0.52% ± 0.22 for the PosVal model on the PosVal dataset and 0.15% ± 0.17 for the NegVal model on the NegVal dataset. Since the gains were rather small, either model shall work well on both positive and negative datasets.

**Figure 5.**
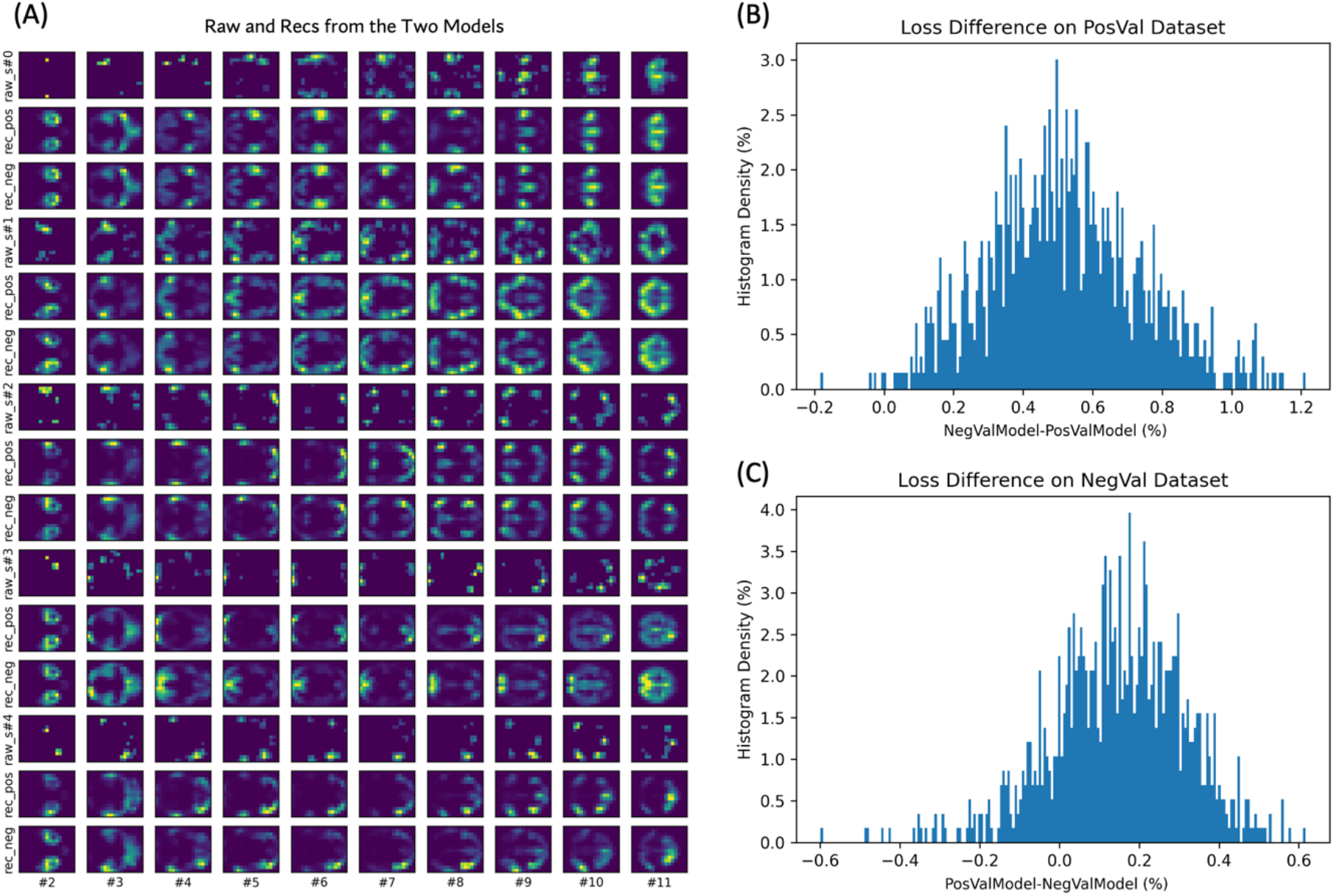
Comparison of PosVal and NegVal models on a separate dataset. (A) The raw and reconstructed images with the two models. Every three rows, in the order of the raw image, the reconstructed image with the model from the PosVal dataset and the reconstructed image with the model from the NegVal dataset, represent one data sample and each column shows one slice of the images. (B-C) Loss differences between the two models on the separate PosVal dataset (B) and NegVal dataset (C), respectively. Abbreviations: PosVal, positive-value; NegVal, negative-value.

### Prediction of transient CAPs with prior latent time courses

To evaluate whether the encoder and the mapping captured the underlying brain dynamics, we predicted the latent variable of the subsequent frame using a segment of latent time courses (40 frames) with the AWD-LSTM model. Examples of comparison between the ground truth latent variables and the predicted ones are depicted in Figure 6; while the comparisons between the ground truth transient CAPs and the transient CAPs of the predicted latent variables are shown in Figure 7.

**Figure 6.**
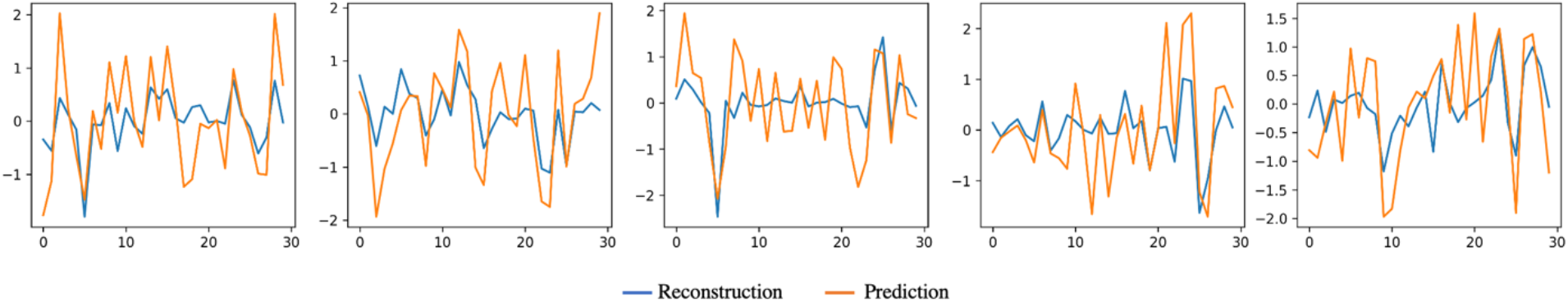
Five random examples of comparison between the reconstructed ground truth latent variables (in blue) and the predicted latent variables (in orange).

**Figure 7.**
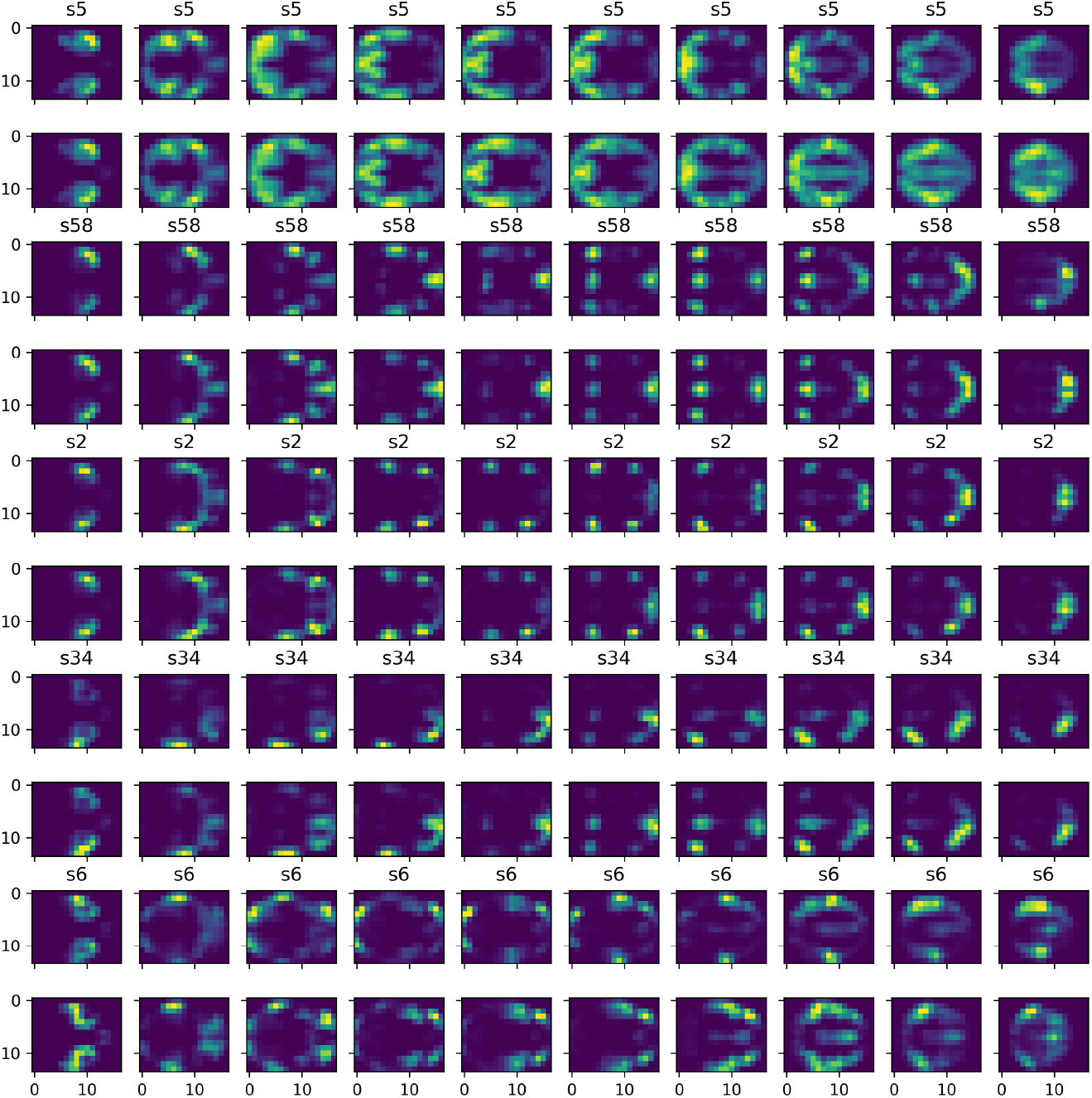
Prediction examples for five randomly selected transient CAPs (s5, s58, s2, s34, and s6). For each case, the upper row shows the reconstructed transient CAP 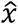 given ground truth *x*, while the lower row shows the reconstructed transient CAP of the predicted latent variable given the 40 frames of transient CAPs prior to *x*.

Despite some variations from the ground truth latent variables, the predicted ones generally matched the ground truth, as seen in Figure 6. The match between ground truth and prediction was closer from the CAPs perspective, as illustrated in Figure 7 where the transient CAPs of predicted latent variables were very similar to the ground truth transient CAPs. Note that neither multi-layer LSTM nor bidirectional LSTM improved the prediction performance, indicating a single-layer forward LSTM model would be sufficient for the temporal modeling.

## DISCUSSION

The present study investigated the representation of framewise transient CAPs in a low-dimension latent space using a generative variational autoencoder neural network(Kingma & Welling 2013). This method was applied on multiple datasets of both low and high resolutions and with both positive and negative values. With LSTM (Hochreiter & Schmidhuber 1997), we tested whether the encoding was effective and carry information of underlying brain dynamics by predicting the transient CAP of brain in the subsequent frame. Our results demonstrated that VAE could embed transient CAPs in latent space in an effective and robust manner and that the encoded latent time courses carried critical brain dynamic information. This provides a new vehicle for modeling the complex spatiotemporal dynamics of brain in a framewise manner.

### The VAE representation

The encoder *q_θ_* (*z/x*) and the decoder *p*_ø_(*x/z*) of VAE essentially provide mappings between the high-dimensional image space and the low-dimensional latent space. Thanks to the powerful deep neural networks, arbitrary complex mappings are trainable through a few simple linear and non-linear activation layers according to the universal approximation (Hornik et al 1989). The trained mappings are only dependent on the two spaces, irrespective of individual subjects. In our study, both patients and controls were used in the model training and the trained models worked on both groups. Further, the trained model on one dataset (e.g., on B-SNIP) is applicable to other datasets (e.g., ABIDE-I) if the imaging spaces are the same, as shown in Figure 1C. However, differences in the imaging spaces caused by hardware, imaging protocols, atlas selections and other factors usually call for separate model training for fine mappings.

Another advantage of VAE is that the samples in the latent space follow the same multivariate Gaussian distribution. This adds critical constraints on the complex mapping that could be arbitrary and ensures the mapping smoothness. Therefore, minor variations in the latent variables did not change the transient CAPs in the image space significantly, as illustrated in Figures 6 and 7.

### Positive and negative VAE models

The present study trained and compared the VAE models using both positive values and negative values of normalized rs-fMRI data. We found the two models were very similar, suggesting that the negative-value frames reflect similar transient brain dynamic patterns as the positive-value ones do. This is in line with the theory of dynamic functional organization of brain supported by the same structural connectome (Smith et al 2009).

The benefit of this finding is that the negative-value time frames, which were usually discarded in prior studies, can provide additional training samples for neural network modelling of framewise brain dynamic analysis. In our settings, thresholded transient CAPs by the 15^th^ and the 85^th^ percentiles simply double the train samples in the original analysis.

### Prediction of transient CAPs via AWD-LSTM

With the constructed latent time courses, we were able to predict the transient brain CAP in the subsequent future frame based on the preceding 40 frames using AWD-LSTM (Merity et al 2017). Compared with the ground-truth transient CAPs, the predictions were fairly accurate, despite minor differences in details, indicating the AWD-LSTM algorithm could effectively model the relationship between the current frame and the prior ones. This is in line with Sobczak and colleagues’ work where a gated recurrent unit RNN model successfully predicted future temporal evolution of rs-fMRI data (Sobczak et al 2021). Nevertheless, it is possible that the high prediction accuracy was mostly a result of the autocorrelation amongst nearby frames, rather than the modeling power of the recurrent neural network. If so, the prediction accuracy for the transient CAPs of the next frame might be compromised if there are changes in the brain states. This is an issue that warrants further study.

Another issue to note is that the prediction model is dependent on the time interval (TR) between frames. This brings challenges to train the prediction model on heterogenous datasets with different TRs like ABIDE or to transfer a trained model to a new dataset with different TR. One possible solution might be temporal resampling so that the resampled frames have the same time intervals across datasets.

## CONCLUSION

With the variational autoencoder neural network, transient co-activity patterns of fMRI images, either co-activated or co-deactivated, can be effectively represented in a low-dimensional latent space, where the complex dynamics of brain activity could be successfully modeled in a framewise manner via LSTM recurrent neural network.

## ACKNOWLEDGEMENT

The authors would like to thank HCP, ABIDE and B-SNIP projects for the data sharing. These datasets can be accessed at: 1) HCP (https://www.humanconnectome.org/study/hcp-young-adult/document/1200-subjects-data-release); 2) ABIDE-1 (http://fcon 1000.projects.nitrc.org/indi/abide/): and 3) B-SNIP (https://nda.nih.gov/edit_collection.html?id=2274).

